# Bioinformatic analysis of differentially expressed long non-coding RNAs in skeletal muscle following aerobic and resistance exercise

**DOI:** 10.1101/2025.10.20.683427

**Authors:** Kassia Régnier, Lucas P.R. Beaupre, Ian F. Coccimiglio, Taylor J. McColl, David C. Clarke, Brendon J. Gurd

**Author notes:** Corresponding author and lead contact: Brendon J. Gurd.

## Abstract

Emerging evidence suggests that long non-coding RNA (lncRNA) molecules influence the adaptive response to exercise, but how lncRNA responses differ between endurance and resistance exercise modalities is poorly understood. The purpose of this study was to bioinformatically infer the expression of lncRNA in skeletal muscle following acute aerobic exercise (AE) and resistance exercise (RE). We downloaded publicly available RNA-seq data, performed a differential expression analysis, and compared lncRNA expression profiles between different exercise types (AE vs. RE) at three time points: baseline, 1 hour post-exercise, and 4 hours post-exercise. We observed distinct lncRNA profiles between acute AE and RE at different time points, suggesting that lncRNA perform different roles in controlling the response to different exercise modalities in skeletal muscle. Future studies should investigate the specific roles of these lncRNAs in the response to acute exercise in skeletal muscle.

## Introduction

Skeletal muscle exhibits remarkable plasticity, responding and adapting to diverse stimuli (1). Exercise induces molecular responses in skeletal muscle at the nucleic acid and protein levels (2) with the nature and magnitude of response dependent on modality, duration, and intensity. Broadly speaking, both aerobic exercise (AE) and resistance exercise (RE) stimulate muscle protein synthesis (MPS), but the specific proteins synthesized differ, as do the subsequent phenotypic adaptations (3). AE induces mitochondrial biogenesis, leading to increased oxidative capacity and resistance to fatigue (4), while RE results in increased actin and myosin content, muscle fiber cross-sectional area, and strength (4–6). The different phenotypes elicited by AE and RE are attributed to the specific molecular responses elicited by each type of exercise – responses typically characterized by observing changes in intracellular signaling, mRNA expression, and protein content.

Over the past decade, long-noncoding RNA molecules (lncRNA), which are RNA molecules longer than 200 nucleotides that do not encode proteins [7–10], have been implicated in many biological processes including the adaptive response to exercise (9,11,12). LncRNAs are thought to function primarily as regulators of gene expression through epigenetic, transcriptional, and non-transcriptional mechanisms (13). For example, the lncRNA Xist is involved in X-chromosome silencing through chromosome coating and the recruitment of epigenetic regulators (14). LncRNAs can also interact with mRNA and microRNA (miRNA) to regulate diverse cellular processes including gene expression, cellular development, chromatin regulation, and gene splicing (13).

Several lncRNA appear to contribute to the regulation of the adaptive response to exercise in skeletal muscle. The lncRNA CYTOR, for example, can modify chromatin accessibility of target genes (10). CYTOR expression declines with age, and is implicated in age-related loss of muscle mass and strength (10). The lncRNA Tug1 decreases PPARGC1A expression and impairs mitochondrial function, when knocked down in mouse myotubes (15). Further, in response to treadmill running in mice, Guo et al. (2022), identified VEGF, mTOR, and NF-kB as potential targets of lncRNA regulation (16). Despite this preliminary evidence, our understanding of the exercise mediated lncRNA response in skeletal muscle remains in its infancy, with the majority of available evidence being gleaned from animal, and cellular models.

In human skeletal muscle, initial evidence suggests that different modalities of exercise training induce distinct lncRNA expression patterns (9). Specifically, twelve weeks of high-intensity interval training (HIIT), resistance training (RT), and combined RT and HIIT induced differential expression of 204, 43, and 15 lncRNAs, respectively, while eight weeks of endurance training caused differential expression in 52 lncRNAs (9). Few lncRNA were differentially expressed in response to more than one training type, suggesting specificity of lncRNA responses. Although these results shed light on the adaptive response to training, the effects of acute AE and RE on lncRNA expression remain unstudied. Based on the lncRNA response to exercise training (9), it is reasonable to hypothesize that acute exercise will serve as a stimulus for lncRNA expression in skeletal muscle with expression profiles being specific to exercise modality (i.e. AE vs RE) specific.

The purpose of this study was to identify differentially expressed lncRNA following acute AE and RE, and to test the hypothesis that distinct lncRNA expression profiles exist between exercise modalities. To assess this hypothesis, we analyzed a publicly available RNA-seq dataset [GSE107934, (17)] of skeletal muscle gene expression measured before, 1 hour, and 4 hours post-exercise. Testing this hypothesis will add to the broader knowledge of the molecular response to acute exercise and provide further evidence regarding the lncRNA response to acute exercise in human skeletal muscle. Further, our study results will add to the currently limited knowledge pertaining to lncRNAs and exercise in humans.

## Methods

### Study Design

The current study is a retrospective analysis of publicly deposited RNA-seq data collected by Dickinson et al. (2018). The ethical approval for the study was obtained through Midwestern University. Dickinson et al. (2018) [GSE107934, (17)] compared transcriptomic responses between aerobic and resistance exercise using RNA-seq. The experiment involved six recreationally active healthy young men who did not participate in regular exercise training programs of more than two days per week (means□±□SD: 27□±□3 yr, 179□±□6 cm, 79□±□10 kg).

Full experimental details are presented in Dickinson et al. (2018). Briefly, participants visited the laboratory twice, once to perform AE and the other RE. The trials were separated by approximately one week, and the exercise types were assigned in a randomized, counterbalanced crossover fashion. In the first exercise trial, a skeletal muscle biopsy was obtained following 30 minutes of supine rest and was used as a control sample for the two exercise trials. Each exercise bout featured a 5-minute warm up at a low workload on a stationary bike. The AE bout was comprised of 40 minutes of stationary cycling at ∼70% maximal heart rate. The RE bout consisted of eight sets of 10 repetitions of isotonic unilateral leg extensions with each leg, at ∼60-65% of participant 1-repetition maximum, with three minutes of rest between each set. The RE bout also lasted approximately 40 minutes. All biopsies were obtained from the vastus lateralis: control samples from the non-dominant leg, post-first exercise sample from the dominant leg, and post-second exercise sample from the same leg as the control biopsy. Following the RE bout, the biopsied leg was the first to perform the unilateral exercise. Immediately upon collection of the muscle tissue, samples were blotted, connective and adipose tissue were dissected, and samples were frozen in liquid nitrogen and stored at -80 °C until analysis (17). All participants provided written, informed consent prior to participating in the study.

RNA was isolated from frozen muscle tissue and sequenced on an Illumina HiSeq 2500 instrument using 75-bp paired-end reads (Illumina, San Diego, CA). The average sequencing depth across samples was 24.3 million reads. One sample from the AE at 4 hours post-exercise failed library preparation and was therefore eliminated from the analysis. FastQC (18) was used to monitor read quality of the RNA-seq data. Principal component analysis (PCA) was performed to determine any outlier samples using a cutoff of 1.5× outside the interquartile range (no samples were deemed outliers).

### Methodological Approach to the Problem

We used RStudio to perform the bioinformatic analysis in this study (v.1.4.1103, [24]), which followed a modified version of the workflow *Analyzing RNA-seq data with DESeq2* (v.1.30.1, [25,26]):

https://www.bioconductor.org/packages/devel/workflows/vignettes/rnaseqGene/inst/doc/rnaseqGene.html#pca-plot.

Specifically, the ExpressionSet object (containing the experiment data) was downloaded using the GEOquery package (v.2.58.0, (22)). Phenotype data were extracted from the ExpressionSet to create a phenotype table containing participant ID, exercise mode, and biopsy timepoint for each sample.. The phenotype table with the phenotype data was used for the differential expression (DE) analysis. Count data was obtained using the `getGEOSuppFiles` function from GEOquery.

### Differential Gene Expression Analysis

In the DE analysis, five conditions were designated: Baseline, 1 hour post AE, 4 hours post AE, 1 hour post RE, and 4 hours post RE. Control samples (before exercise) were used as the reference level. To measure the effects of exercise mode and time, the five conditions were specified as the design in the DESeqDataSet object. Low-count genes (<1) in the DESeqDataSet were removed from further analysis. The function `DESeq()` from DESeq2 was used to compute the differential expression analysis. Briefly, DESeq() performs the following steps:

1. Size factor estimation (normalization; `estimateSizeFactors`): Normalizes for differences in sequencing depth across samples by computing the median ratio of observed samples (23).
2. Dispersion estimation (`estimateDispersions`): The sum of the biological variance and the shot noise (technical variance). An estimate of the dispersion is reported for each gene (20).
3. Negative binomial generalized linear model (GLM) fitting and Wald statistics (`nbinomWaldTest`): Evaluates the significance of regression coefficients using a negative binomial generalized linear model, which uses the size factors and dispersion estimates (20).

Following the DE analysis, the `resultsNames` function was used to confirm the names of each condition in the design. A boxplot was created to visualize the distribution of the normalized count data. As RNA-seq data is often heteroscedastic, a variance stabilizing transformation (VST) of the count data was applied to provide count values that are approximately homoscedastic. To visualize this transformation, a box plot of the VST data was created. Following the VST, a PCA of the VST data was performed (with time and exercise as groups of interests) to reduce the dimensionality of the VST data, observe trends, and clusters in the data. Subsequently, the Euclidean distance was calculated between samples using the VST data and a heatmap of the sample-to-sample distances was created. Thereafter, the dispersion estimates of the DESeq2 negative binomial model were plotted, with the mean of normalized counts on the x-axis and the dispersion on the y-axis.

We obtained a table of the results (`results`) that displayed the adjusted *P*-value, log fold change (LFC) > 0, LFC < 0, outliers, low counts, and mean count. The `lfcShrink` function (24) was then applied to reduce noise in the estimated log_2_ fold changes, which improves visualization and gene ranking. MA plots were created to visualize gene expression, each MA plot displays the log2-fold changes due to a given variable over the mean of normalized counts for all samples in a DESeqDataSet. Finally, the results obtained from results (“Normal”), were plotted followed by the shrunken results.

### Filtering Results

The results were filtered using a custom function according to the criteria in Dickinson et al. (2018): adjusted *P*-values less than 0.05 and absolute log2 fold change values less than(?) 0.58. Results were annotated using a custom function based on Ensembl gene IDs. The biomaRt package (v.2.49.2, (25)) was used to retrieve Ensembl gene ID’s, HGNC gene symbols, gene biotype, chromosome name, gene start and end positions, strand, and gene description according to the GRCh37.p13 assembly (consistent with Dickinson et al. (2018)). The two created functions were applied to the results of each coefficient. Finally, the results were exported into individual csv files, where lncRNAs were filtered from other genes and transcripts (according to the ‘gene_biotype’ classification).

### Validation of Results

Differential expression results were validated against the results reported in Dickinson et al. (2018). We extracted the results from their supplemental files, which featured the following comparisons: baseline vs. post-exercise, DE genes following AE vs. RE, unique DE in each exercise mode by time, unique DE for AE at each timepoint, and unique DE for RE at each timepoint. The results were compared using base R functions and functions from the stringi package (v.1.7.4, (26)). Finally, each comparison from R was exported into individual csv files.

### MetaMex

We further examined lncRNAs expression in response to AE and RE using the MetaMex database (https://www.metamex.eu) (27). We searched MetaMex for each lncRNA we identified from the Dickinson et al. (2018) dataset. Our search criteria included acute aerobic and resistance exercise, with the following search parameters: (i) timepoints: all post-exercise timepoints (immediate to 96 hours); (ii) sex: male, female, or undefined; (iii) age: young, middle-aged, and elderly participants; (iv) fitness: sedentary, active, and athlete populations; (v) weight: lean, overweight, obese, and class III obese participants; (vi) muscle tissue: vastus lateralis, biceps brachii, soleus, and gastrocnemius; (vii) health status: healthy, impaired glucose tolerance, type 1 and type 2 diabetes, metabolic syndrome, chronic kidney disease, peripheral arterial disease, chronic obstructive pulmonary disease, Parkinson’s disease, chronic heart failure, frailty, and sarcopenia.

## Results

### Validation of Findings

Our analysis revealed 40 DE genes unique to AE (36 were the same as found by Dickinson et al. [2018], who reported a total of 48), and 295 DE genes unique to RE (280 were the same as found by Dickinson et al. [2018], who reported a total of 348). One hour post AE, 40 DE genes were observed (Dickinson et al. [2018] reported a total of 48), 4 hours post AE, 204 DE genes were observed (Dickinson et al. [2018] reported a total of 221). One hour post RE, 60 DE genes were observed (Dickinson et al. [2018] reported a total of 67), 4 hours post RE, 474 DE genes were observed (Dickinson et al. [2018], reported a total of 523).

### Principal Component Analysis

The PCA of the VST data revealed sample-like patterns in gene expression levels between groups (Figure 1). The first principal component (PC) explained 41% of the variance, while the second PC explained 15% of the variance. The PCA confirmed the distinct expression pattern of the samples according to the biopsy timepoint, however, the expression pattern for different exercise types was not as clear.

**Figure 1.**
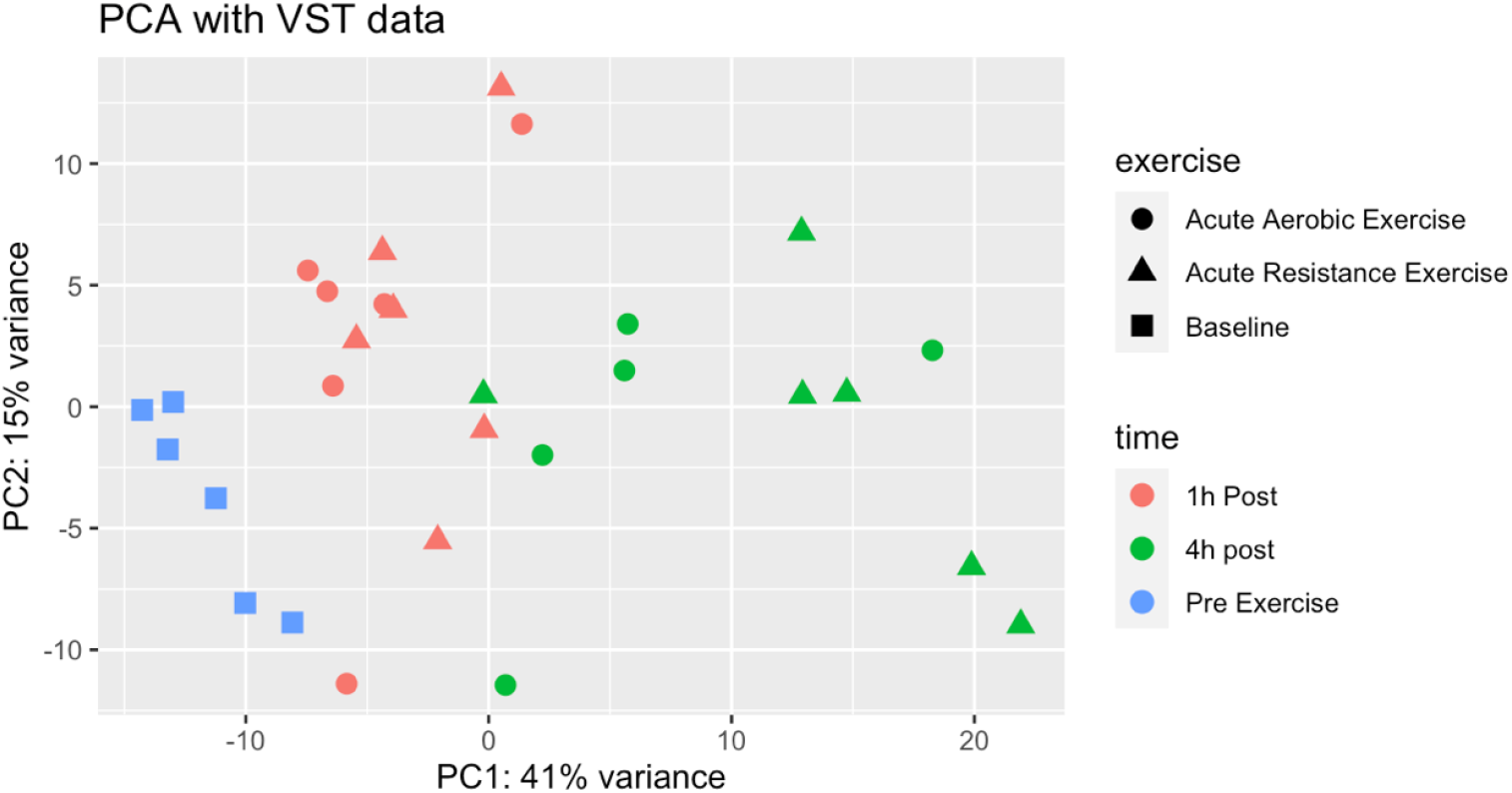
PCA plot of the variance stabilized transformed data with exercise and time as interest groups. The plot shows the first two components generated by the PCA. Each point on the plot represents one sample. Samples are identified based on exercise, and time and are labeled in the legend. The greater the distance between the points, the greater the difference of gene expression patterns. The distance between the points between different conditions is proportional to their variance.

### Differential Expression of LncRNAs

A total of 27 lncRNAs were differentially expressed among both exercise types (Figure 2). Among these, 20 lncRNAs were upregulated, while seven lncRNAs were downregulated (Table 1). DE of lncRNA’s in response to AE and RE at 1 hour and 4 hours post exercise are presented in Figure 2. Notably, lncRNA are temporally expressed, mostly 4 hours after both AE and RE. RE also appears to be a stimulus for a larger amount of lncRNAs.

**Table 1.**
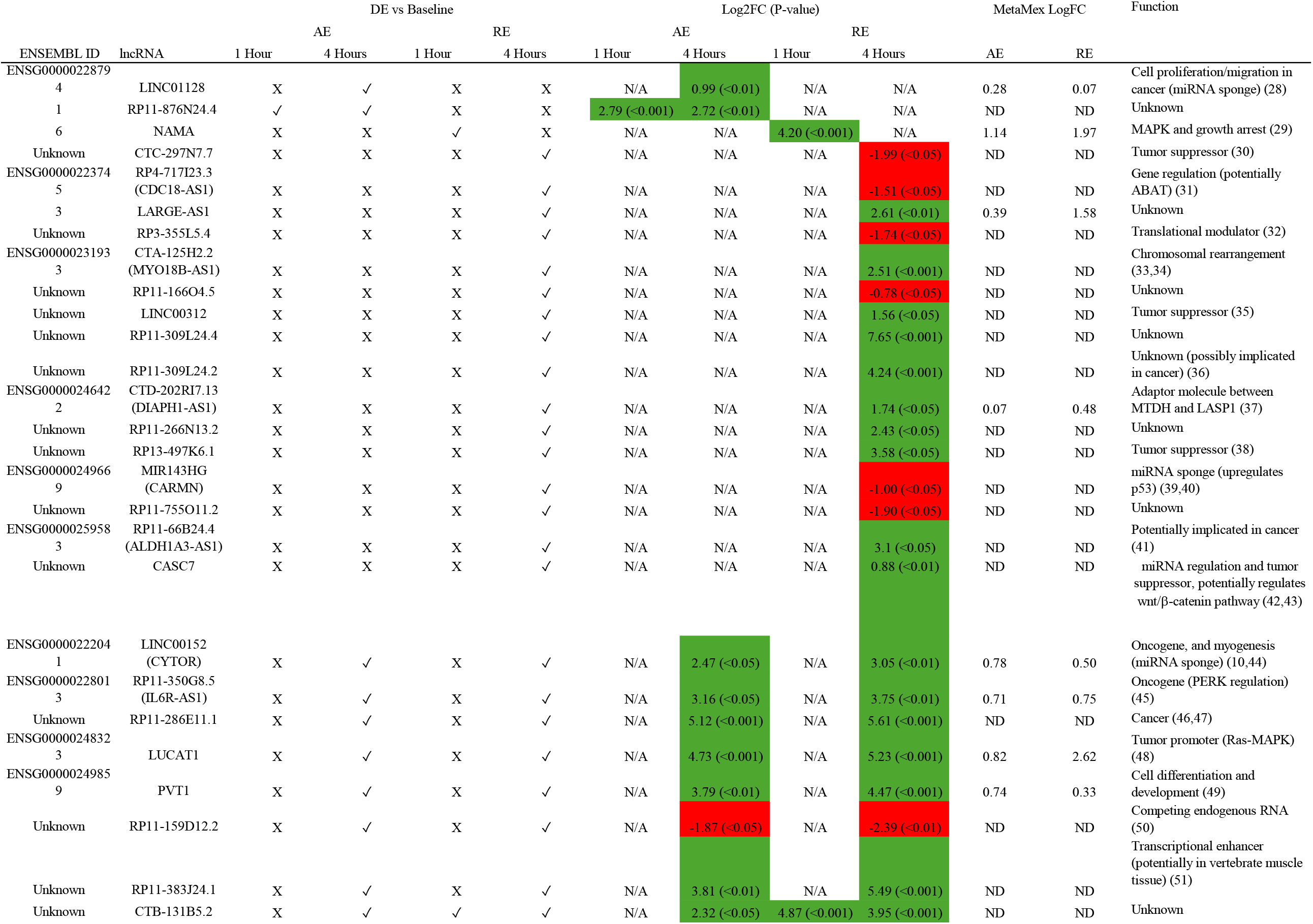
Identification, and quantified expression of lncRNAs that are differentially expressed following exercise.

**Figure 2.**
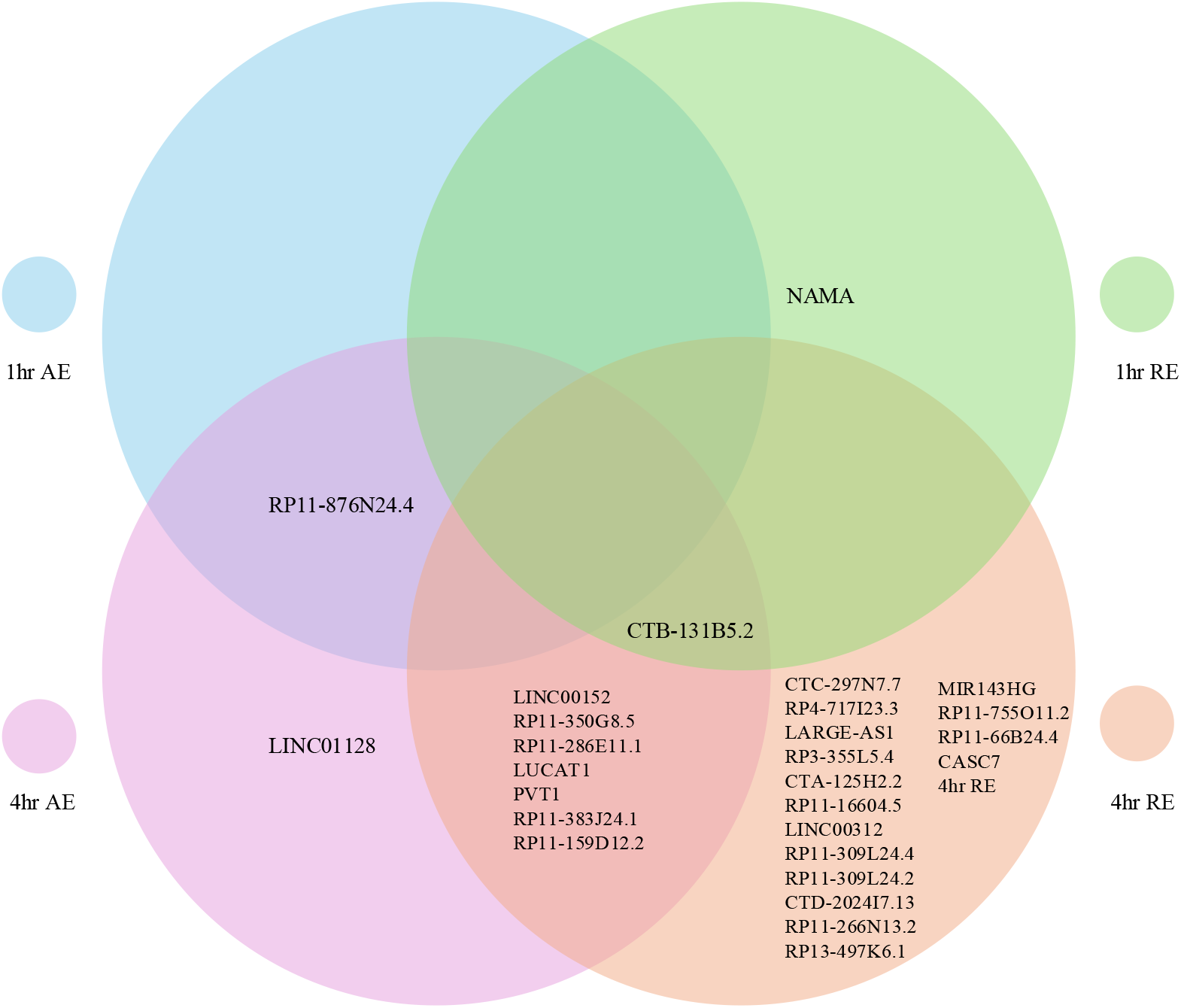
LncRNAs identified1 hour, and 4 hours post AE, and 1 hour, and 4 hours post RE.

### MetaMex

Eight of the DE lncRNAs that we identified appeared in the MetaMex search (Table 1). Further, three lncRNAs were reported to be DE following AE in the MetaMex database but were not differentially expressed in our analysis. However, these same three lncRNAs were DE after RE in in both our analysis and the MetaMex data.

## Discussion

The current study used publicly available RNA-seq data (17) to investigate the expression profiles of lncRNAs in skeletal muscle following acute AE and RE at two post-exercise timepoints. To characterize lncRNA expression patterns following different types of acute exercise, we compared differentially expressed lncRNAs between exercise types (AE vs. RE) across different time points (baseline, 1 hour post-exercise, 4 hours post-exercise). For the first time, we report that lncRNAs are differentially expressed predominantly four hours post-exercise, and only slightly differentially expressed one-hour post-exercise. These findings emphasize the unique transcriptomic responses of skeletal muscle to different acute exercise modalities and highlight the potentially important role of lncRNAs in mediating the adaptive response to exercise.

### LncRNA Expression Post-Exercise

Molecular expression patterns can reveal molecular signatures associated with specific outcomes (52–54), offering insights into responses to different types of acute exercise. However, most studies investigating gene expression patterns following exercise have focused on mRNA expression. Examining lncRNAs across exercise modalities provides a novel perspective on the skeletal muscle molecular response.

We identified distinct lncRNA expression profiles between acute aerobic and resistance exercise, suggesting that lncRNAs might have unique roles in the skeletal muscle adaptive response to acute exercise. In total, 20 lncRNAs were upregulated, while seven lncRNAs were downregulated; 10 lncRNAs were differentially expressed following AE, and 25 after RE, only eight lncRNAs overlapped between modalities. These findings provide additional insight into the molecular mechanisms that underlie the different phenotypic responses to AE and RE, and suggest that specific lncRNA signatures may contribute to the divergent adaptations to AE and RE (53,55).

The majority of the differentially expressed lncRNAs we identified have yet to be functionally characterized. However, two hypotheses provide clues as to the possible functions of lncRNAs: 1) transcriptional regulation (reviewed in (13,56)) and 2) competitive endogenous RNA (ceRNA) theory (57,58). Our results indicate that LINC00152, also known as the cytoskeleton regulator RNA (CYTOR), was upregulated at 4 hours post exercise in both AE (Log2FC = 2.47) and RE (Log2FC = 3.05). In accordance with the first hypothesis, CYTOR transcriptionally controls gene expression (10). CYTOR is conserved in humans, mice, and rats, and is highly responsive to RE, and to a lesser degree, AE (10). CYTOR promotes skeletal muscle hypertrophy by stimulating muscle progenitor differentiation (10), and exerts this effect by binding and sequestering the transcription factor Tead1, reducing the chromatin accessibility and occupancy of Tead1-binding motifs in its target genes (10). Further, mechanistic evidence suggests Tead1 expression promotes the expression of type I muscle fibres and thus reduces the type II muscle fiber phenotype (59,60). Therefore, CYTOR-mediated suppression of Tead1 expression promotes type II muscle fibers (10). CYTOR knockout models exhibit sarcopenic features, and there is an age-related reduction in CYTOR expression in both rat and human muscle cells (10). However, overexpression of CYTOR in older mice restored muscle mass, fiber size, and reversed the age-related loss of type II muscle fibers (10). It is plausible that lncRNAs identified in our study may function like CYTOR and transcriptionally control gene expression. Future research could test this hypothesis by employing methods such as gain- or loss-of-function experiments to examine how altering specific lncRNA expression alters gene expression relevant in the exercise response.

We found the lncRNA plasmacytoma variant translocation 1 (PVT1) to be DE 4 hours post AE (Log 2FC = 3.79) and RE (Log2FC = 4.47); these results agree with those from MetaMEX (27). Currently, one study has been conducted that determined that PVT1, along with four other lncRNAs is involved in muscle development, and atrophy through regulatory networks involving mRNA, and microRNA(61). Further, evidence suggests PVT1 is implicated in mitochondrial dysfunction in podocytes (62) and has been proposed as a potential biomarker for sarcopenia in human plasma serum (63). In humans, increased PVT1 expression in podocytes was associated with increased mitochondrial dysfunction, while its knock out in mouse models restored mitochondrial function, potentially through AMPK□ ubiquitination (62). Furthermore, PVT1 levels in plasma serum associated with age in humans and with sarcopenia in older adults (63). Notably, six months of aerobic and resistance training in sarcopenic adults decreased PVT1 expression compared with a sarcopenic control group (63). In addition to these functions, the possibility exists that PVT1 may function in the regulation of skeletal muscle via mechanisms that have yet to be elucidated. Based on evidence in podocytes (62), and plasma serum (63), it is reasonable to postulate that PVT1 may regulate mitochondrial function, and muscle development in skeletal muscle. Thus, the mechanisms of PVT1 provide appealing targets for future studies, moreover further research is required to investigate how PVT1 expression changes after exercise.

While our study explored the differential expression of lncRNA at various post-exercise time points across different exercise modalities, the functions of many of these lncRNAs remain unknown. Future studies should emphasize constructing lncRNA-miRNA networks that underlie the response to acute AE and RE to gain further mechanistic insight on the molecular interactions regulating the response to acute exercise. Further, lncRNAs hypothesized to be involved in transcriptional regulation often exert their effects directly on the chromatin, either with a protein intermediate (64), or through the formation of triple helices via hybridization (65,66). Therefore, experiments employing methodologies such as chromatin isolation by RNA purification, RNA antisense purification, and capture hybridization analysis of RNA targets will be increasingly important in understanding the mechanisms of lncRNAs in the response to acute exercise.

### Limitations and Future Directions

Our study features the following noteworthy limitations, several arising from the original dataset on which the analysis is based. First, the study was a quasi-experimental cross-over design featuring a relatively small sample size of young, healthy adult males. The quasi-experimental design limits the ability to make firm causal conclusions. The one-week interval between exercise bouts may have been insufficient for a full washout of effects from the first bout prior to the second. This limitation is further compounded by the use of a single baseline muscle biopsy, obtained prior to the first exercise bout, as the comparator for both exercise bouts. Thus, the true baseline state prior to the second bout of exercise was not captured. Only six participants completed the exercise trials, and one of the 30 samples could not be included, leaving 29 samples in the dataset. Additionally, the lack of female participants means that these findings cannot be generalized to females, which is unfortunate given the known morphological, and physiological differences that exist between sexes, and the underrepresentation of females in exercise science (67). An additional limitation of the study design is the relatively few time points assessed. The response to acute exercise persists beyond 4 hours (68–70), such that future studies should focus on a broader range of time points.

Additional limitations arise from the RNA-seq data. First, lncRNAs tend to be expressed at lower levels than protein-coding genes. Since we used the same cut offs as the original authors, the possibility exists that biologically relevant but marginally expressed lncRNAs were excluded. Another potential limitation of the RNA-seq data is coverage bias, which is the uneven distribution of reads across genomic regions that can lead to inaccurate estimates of expression levels and their differences (71). Coverage bias can arise from differences in transcript size, guanine-cytosine content, sequence complexity, and secondary structure (72,73). In the current study, the average sequencing depth was 24.3 million reads. To accurately detect lncRNAs and other marginally expressed transcripts, higher sequencing depths starting at 100 million reads per sample is recommended (74). Moreover, Dickinson et al. (2018) used a read length of 75 nucleotides in their RNA-seq workflow, however the ideal read length to adequately capture lncRNAs is closer to 100 nucleotides (75). Further, Dickinson et al. (2018) used Poly(A) selection in the library preparation step of their RNA-seq dataset, which enriches mRNAs through hybridization. Many lncRNAs lack PolyA tails, which may cause lncRNAs to be missed in the Poly(A) selection step (76). Superior methods exist for preparing RNA-seq libraries that better target lncRNAs, such as rRNA depletion (77). Therefore, future studies should judiciously choose their library preparation methods depending on the extent to which lncRNA detection is prioritized.

## Conclusion

Our findings highlight the utility of bioinformatics in exercise science, by analyzing a publicly available RNA-seq data to reveal distinct lncRNA expression profiles at different time points following acute AE and RE. We identified 13 DE lncRNAs following AE and 29 DE lncRNAs following RE, reflecting the unique molecular responses to each exercise modality. Further, we proposed potential mechanisms by which lncRNAs contribute to the acute exercise response in skeletal muscle. Characterizing these exercise-specific lncRNA profiles in skeletal muscle represents a key step towards understanding the molecular response to exercise and its potential therapeutic value. Our study provides a framework for future investigations into the mechanistic functions of lncRNAs in relation to exercise. We hope that our study encourages wider use of bioinformatics approaches to analyze omics data and further investigation of lncRNAs in exercise science.

## Author statements

### Author Contributions

Conceptualization: KR, DCC, BJG

Data curation: KR, IFC, TJM

Formal analysis: KR, IFC, TJM

Investigation: KR, IFC, TJM

Methodology: KR, IFC, TJM

Project administration: KR, IFC, TJM, DCC, BJG

Resources: DCC, BJG

Software: KR, IFC, TJM

Supervision: DCC, BJG

Validation: KR, IFC, TJM

Visualization: KR, LRPB, IFC, TJM

Writing – original draft: LPRB, BJG

Writing – review & editing: KR, TJM, DCC

### Data availability statement

Data generated or analyzed during this study are available from the corresponding author upon reasonable request.

## References

1. Ferraro E, Giammarioli AM, Chiandotto S, Spoletini I, Rosano G. Exercise-induced skeletal muscle remodeling and metabolic adaptation: redox signaling and role of autophagy. Antioxid Redox Signal. 2014 July 1;21(1):154–76.

2. Moberg M, Lindholm ME, Reitzner SM, Ekblom B, Sundberg CJ, Psilander N. Exercise Induces Different Molecular Responses in Trained and Untrained Human Muscle. Medicine & Science in Sports & Exercise. 2020 Aug;52(8):1679–90.

3. Egan B, Zierath JR. Exercise Metabolism and the Molecular Regulation of Skeletal Muscle Adaptation. Cell Metabolism. 2013 Feb;17(2):162–84.

4. Scarpulla RC, Vega RB, Kelly DP. Transcriptional integration of mitochondrial biogenesis. Trends Endocrinol Metab. 2012 Sept;23(9):459–66.

5. Dreyer HC, Fujita S, Cadenas JG, Chinkes DL, Volpi E, Rasmussen BB. Resistance exercise increases AMPK activity and reduces 4E□BP1 phosphorylation and protein synthesis in human skeletal muscle. The Journal of Physiology. 2006 Oct 15;576(2):613–24.

6. Terzis G, Georgiadis G, Stratakos G, Vogiatzis I, Kavouras S, Manta P, et al. Resistance exercise-induced increase in muscle mass correlates with p70S6 kinase phosphorylation in human subjects. Eur J Appl Physiol. 2007 Nov 15;102(2):145–52.

7. Cabili MN, Trapnell C, Goff L, Koziol M, Tazon-Vega B, Regev A, et al. Integrative annotation of human large intergenic noncoding RNAs reveals global properties and specific subclasses. Genes Dev. 2011 Sept 15;25(18):1915–27.

8. Derrien T, Johnson R, Bussotti G, Tanzer A, Djebali S, Tilgner H, et al. The GENCODE v7 catalog of human long noncoding RNAs: Analysis of their gene structure, evolution, and expression. Genome Res. 2012 Sept;22(9):1775–89.

9. Bonilauri B, Dallagiovanna B. Long Non-coding RNAs Are Differentially Expressed After Different Exercise Training Programs. Front Physiol. 2020 Sept 15;11:567614.

10. Wohlwend M, Laurila PP, Williams K, Romani M, Lima T, Pattawaran P, et al. The exercise-induced long noncoding RNA CYTOR promotes fast-twitch myogenesis in aging. Sci Transl Med. 2021 Dec 8;13(623):eabc7367.

11. Mattick JS, Amaral PP, Carninci P, Carpenter S, Chang HY, Chen LL, et al. Long non-coding RNAs: definitions, functions, challenges and recommendations. Nat Rev Mol Cell Biol. 2023 June;24(6):430–47.

12. Li G, Deng L, Huang N, Sun F. The Biological Roles of lncRNAs and Future Prospects in Clinical Application. Diseases. 2021 Jan 13;9(1):8.

13. Statello L, Guo CJ, Chen LL, Huarte M. Gene regulation by long non-coding RNAs and its biological functions. Nat Rev Mol Cell Biol. 2021 Feb;22(2):96–118.

14. Li J, Ming Z, Yang L, Wang T, Liu G, Ma Q. Long noncoding RNA XIST: Mechanisms for X chromosome inactivation, roles in sex-biased diseases, and therapeutic opportunities. Genes & Diseases. 2022 Nov;9(6):1478–92.

15. Trewin AJ, Silver J, Dillon HT, Della Gatta PA, Parker L, Hiam DS, et al. Long non-coding RNA Tug1 modulates mitochondrial and myogenic responses to exercise in skeletal muscle. BMC Biol. 2022 Dec;20(1):164.

16. Guo J, Yuan Y, Zhang L, Wang M, Tong X, Liu L, et al. Effects of exercise on the expression of long non□coding RNAs in the bone of mice with osteoporosis. Exp Ther Med. 2021 Nov 23;23(1):70.

17. Dickinson JM, D’Lugos AC, Naymik MA, Siniard AL, Wolfe AJ, Curtis DR, et al. Transcriptome response of human skeletal muscle to divergent exercise stimuli. Journal of Applied Physiology. 2018 June 1;124(6):1529–40.

18. Andrews S. FastQC: A Quality Control tool for High Throughput Sequence Data. Babraham Institute [Internet]. 2010 [cited 2025 July 10]; Available from: https://www.bioinformatics.babraham.ac.uk/projects/fastqc/

19. Posit Team. RStudio: Integrated Development Environment for R [Internet]. PBC; 2025. Available from: http://www.posit.co/

20. Love MI, Huber W, Anders S. Moderated estimation of fold change and dispersion for RNA-seq data with DESeq2. Genome Biol [Internet]. 2014 Dec 5 [cited 2025 July 10];15(12). Available from: https://genomebiology.biomedcentral.com/articles/10.1186/s13059-014-0550-8

21. Love MI, Anders S, Kim V, Huber W. RNA-Seq workflow: gene-level exploratory analysis and differential expression. F1000Res. 2015 Oct 14;4:1070.

22. Davis S, Meltzer PS. GEOquery: a bridge between the Gene Expression Omnibus (GEO) and BioConductor. Bioinformatics. 2007 July 15;23(14):1846–7.

23. Anders S, Huber W. Differential expression analysis for sequence count data. Genome Biol [Internet]. 2010 Oct [cited 2025 July 10];11(10). Available from: https://genomebiology.biomedcentral.com/articles/10.1186/gb-2010-11-10-r106

24. Zhu A, Ibrahim JG, Love MI. Heavy-tailed prior distributions for sequence count data: removing the noise and preserving large differences. Stegle O, editor. Bioinformatics. 2019 June 1;35(12):2084–92.

25. Durinck S, Spellman PT, Birney E, Huber W. Mapping identifiers for the integration of genomic datasets with the R/Bioconductor package biomaRt. Nat Protoc. 2009 Aug;4(8):1184–91.

26. Gagolewski M. stringi: Fast and Portable Character String Processing in R. J Stat Soft [Internet]. 2022 [cited 2025 July 10];103(2). Available from: https://www.jstatsoft.org/v103/i02/

27. Pillon NJ, Gabriel BM, Dollet L, Smith JAB, Sardón Puig L, Botella J, et al. Transcriptomic profiling of skeletal muscle adaptations to exercise and inactivity. Nat Commun. 2020 Jan 24;11(1):470.

28. Zhong M, Fang Z, Ruan B, Xiong J, Li J, Song Z. LINC01128 facilitates the progression of pancreatic cancer through up-regulation of LDHA by targeting miR-561-5p. Cancer Cell Int [Internet]. 2022 Dec [cited 2025 July 7];22(1). Available from: https://cancerci.biomedcentral.com/articles/10.1186/s12935-022-02490-5

29. Yoon H, He H, Nagy R, Davuluri R, Suster S, Schoenberg D, et al. Identification of a novel noncoding RNA gene, NAMA, that is downregulated in papillary thyroid carcinoma with BRAF mutation and associated with growth arrest. Intl Journal of Cancer. 2007 Aug 15;121(4):767–75.

30. Zhu S, Huang X, Zhang K, Tan W, Lin Z, He Q, et al. Low expression of long noncoding RNA CTC-297N7.9 predicts poor prognosis in patients with hepatocellular carcinoma. Cancer Med. 2019 Dec;8(18):7679–92.

31. Chen Y, Zhao G, Li N, Luo Z, Wang X, Gu J. Role of 4□aminobutyrate aminotransferase (ABAT) and the lncRNA co□expression network in the development of myelodysplastic syndrome. Oncol Rep. 2019 Aug;42(2):509–20.

32. Li J, Yuan X, Ma C, Li J, Qu G, Yu B, et al. LncRNA LBX2-AS1 impacts osteosarcoma sensitivity to JQ-1 by sequestering miR-597-3p away from BRD4. Front Oncol [Internet]. 2023 Mar 24 [cited 2025 July 8];13. Available from: https://www.frontiersin.org/articles/10.3389/fonc.2023.1139588/full

33. Toujani S, Tucker EJ, Akloul L, Mary L, Pimentel C, Launay E, et al. Pseudodicentric Chromosome Originating from an X-Autosome Translocation in a Male Patient with Cryptozoospermia. Cytogenet Genome Res. 2022;162(3):124–31.

34. Buckley PG, Mantripragada KK, Benetkiewicz M, Tapia-Páez I, Ståhl TD de, Rosenquist M, et al. A full-coverage, high-resolution human chromosome 22 genomic microarray for clinical and research applications. Human Molecular Genetics. 2002 Dec 1;11(25):3221–9.

35. Zhang R, Jiang Y, Gu J, Zhang X, Xie Y. Diagnostic role of circulating long non-coding RNA LINC00312 in patients with non-small cell lung cancer: a retrospective study. BMC Cancer [Internet]. 2025 Jan 9 [cited 2025 July 8];25(1). Available from: https://bmccancer.biomedcentral.com/articles/10.1186/s12885-024-13393-1

36. Gao Y, Li X, Zhi H, Zhang Y, Wang P, Wang Y, et al. Comprehensive Characterization of Somatic Mutations Impacting lncRNA Expression for Pan-Cancer. Mol Ther Nucleic Acids. 2019 Dec 6;18:66–79.

37. Li ZX, Zheng ZQ, Yang PY, Lin L, Zhou GQ, Lv JW, et al. WTAP-mediated m6A modification of lncRNA DIAPH1-AS1 enhances its stability to facilitate nasopharyngeal carcinoma growth and metastasis. Cell Death Differ. 2022 June;29(6):1137–51.

38. Kalmar A, Nagy ZB, Galamb O, Wichmann B, Bartak BK, Valcz G, et al. PO-377 Whole transcriptome analysis reveals colorectal cancer-associated long non-coding RNAs including UCA1 already dysregulated in colorectal adenomas. ESMO Open. 2018 June;3:A169–70.

39. Wang P, Bao W, Liu X, Xi W. LncRNA miR143HG inhibits the proliferation of glioblastoma cells by sponging miR-504. International Journal of Neuroscience. 2022 Nov 2;132(11):1137–42.

40. Wang X, Wu S, Yang Y, Zhao J. LncRNA CARMN Affects Hepatocellular Carcinoma Prognosis by Regulating the miR□192□5p/LOXL2 Axis. Li G, editor. Oxidative Medicine and Cellular Longevity. 2022 Jan;2022(1):9277360.

41. Fan Y, Cui J, Zhu Q. Heterogeneous graph inference based on similarity network fusion for predicting lncRNA–miRNA interaction. RSC Adv. 2020;10(20):11634–42.

42. Sun W, Yin D. Long noncoding RNA CASC7 inhibits the proliferation and migration of papillary thyroid cancer cells by inhibiting miR-34a-5p. The Journal of Physiological Sciences. 2021;71(1):9.

43. Sun W, Wang D, Zu Y, Deng Y. Long noncoding RNA CASC7 is a novel regulator of glycolysis in oesophageal cancer via a miR-143-3p-mediated HK2 signalling pathway. Cell Death Discov [Internet]. 2022 Apr 26 [cited 2025 July 8];8(1). Available from: https://www.nature.com/articles/s41420-022-01028-y

44. Yue B, Cai D, Liu C, Fang C, Yan D. Linc00152 Functions as a Competing Endogenous RNA to Confer Oxaliplatin Resistance and Holds Prognostic Values in Colon Cancer. Molecular Therapy. 2016 Dec;24(12):2064–77.

45. Grillone K, Ascrizzi S, Cremaschi P, Amato J, Polerà N, Croci O, et al. An unbiased lncRNA dropout CRISPR-Cas9 screen reveals RP11-350G8.5 as a novel therapeutic target for multiple myeloma. Blood. 2024 Oct 17;144(16):1705–21.

46. Zhang P, Xu K, Wang J, Zhang J, Quan H. Identification of N6-methylandenosine related LncRNAs biomarkers associated with the overall survival of osteosarcoma. BMC Cancer. 2021 Dec 1;21(1):1285.

47. Zhang L, Meng X, Zhu X wei, Yang D cheng, Chen R, Jiang Y, et al. Long non-coding RNAs in Oral squamous cell carcinoma: biologic function, mechanisms and clinical implications. Mol Cancer [Internet]. 2019 Dec [cited 2025 July 7];18(1). Available from: https://molecular-cancer.biomedcentral.com/articles/10.1186/s12943-019-1021-3

48. Wu X, Song L, Chen X, Zhang Y, Li S, Tang X. Long non-coding RNA LUCAT1 regulates the RAS pathway to promote the proliferation and invasion of malignant glioma cells through ABCB1. Experimental Cell Research. 2022 Dec;421(2):113390.

49. Wu F, Zhu Y, Zhou C, Gui W, Li H, Lin X. Regulation mechanism and pathogenic role of lncRNA plasmacytoma variant translocation 1 (PVT1) in human diseases. Genes & Diseases. 2023 May;10(3):901–14.

50. Chen L, Zhao T. Identification of KHSRP-Regulated RNAs in Esophageal Cancer by Integrated Bioinformatics Analysis. Cancer Biotherapy and Radiopharmaceuticals. 2021 June 1;36(5):412–24.

51. Bonventre JA, Holman C, Manchanda A, Codding SJ, Chau T, Huegel J, et al. Fer1l6 is essential for the development of vertebrate muscle tissue in zebrafish. Mol Biol Cell. 2019 Feb 1;30(3):293–301.

52. Iyer P, Asante DM, Vyavahare S, Jin LT, Ahluwalia P, Kolhe R, et al. Transcriptional Signatures of Aerobic Exercise-Induced Muscle Adaptations in Humans. J Funct Morphol Kinesiol. 2025 July 19;10(3):281.

53. Drummond MJ, Fujita S, Abe T, Dreyer HC, Volpi E, Rasmussen BB. Human muscle gene expression following resistance exercise and blood flow restriction. Med Sci Sports Exerc. 2008 Apr;40(4):691–8.

54. Furrer R, Handschin C. Molecular aspects of the exercise response and training adaptation in skeletal muscle. Free Radic Biol Med. 2024 Oct;223:53–68.

55. Bori Z, Zhao Z, Koltai E, Fatouros IG, Jamurtas AZ, Douroudos II, et al. The effects of aging, physical training, and a single bout of exercise on mitochondrial protein expression in human skeletal muscle. Exp Gerontol. 2012 June;47(6):417–24.

56. Voorn EL, Koopman FS, Nollet F, Brehm MA. Individualized Aerobic Exercise in Neuromuscular Diseases: A Pilot Study on the Feasibility and Preliminary Effectiveness to Improve Physical Fitness. Physical Therapy [Internet]. 2021 Mar 3 [cited 2025 July 9];101(3). Available from: https://academic.oup.com/ptj/article/doi/10.1093/ptj/pzaa213/6039324

57. Salmena L, Poliseno L, Tay Y, Kats L, Pandolfi PP. A ceRNA Hypothesis: The Rosetta Stone of a Hidden RNA Language? Cell. 2011 Aug;146(3):353–8.

58. Thomson DW, Dinger ME. Endogenous microRNA sponges: evidence and controversy. Nat Rev Genet. 2016 May;17(5):272–83.

59. Tsika RW, Schramm C, Simmer G, Fitzsimons DP, Moss RL, Ji J. Overexpression of TEAD-1 in transgenic mouse striated muscles produces a slower skeletal muscle contractile phenotype. J Biol Chem. 2008 Dec 26;283(52):36154–67.

60. Zhang D, Wang X, Li Y, Zhao L, Lu M, Yao X, et al. Thyroid hormone regulates muscle fiber type conversion via miR-133a1. Journal of Cell Biology. 2014 Dec 22;207(6):753–66.

61. Wenlun W, Chaohang Y, Yan H, Wenbin L, Nanqing Z, Qianmin H, et al. Developing a ceRNA-based lncRNA-miRNA-mRNA regulatory network to uncover roles in skeletal muscle development. Front Bioinform. 2025 Jan 15;4:1494717.

62. Lv Z, Wang Z, Hu J, Su H, Liu B, Lang Y, et al. LncRNA PVT1 induces mitochondrial dysfunction of podocytes via TRIM56 in diabetic kidney disease. Cell Death Dis. 2024 Sept 30;15(9):697.

63. Aparicio P, López-Royo T, Navarrete-Villanueva D, Gómez Cabello AM, González-Gross M, Ara I, et al. Serum lncRNAs NEAT1, PVT1 and H19 as novel biomarkers for sarcopenia diagnosis and treatment response. Non-coding RNA Research. 2025 Oct;14:166–76.

64. Sato M, Kadomatsu T, Miyata K, Warren JS, Tian Z, Zhu S, et al. The lncRNA Caren antagonizes heart failure by inactivating DNA damage response and activating mitochondrial biogenesis. Nat Commun [Internet]. 2021 May 5 [cited 2025 July 10];12(1). Available from: https://www.nature.com/articles/s41467-021-22735-7

65. Schmitz KM, Mayer C, Postepska A, Grummt I. Interaction of noncoding RNA with the rDNA promoter mediates recruitment of DNMT3b and silencing of rRNA genes. Genes Dev. 2010 Oct 15;24(20):2264–9.

66. Martianov I, Ramadass A, Serra Barros A, Chow N, Akoulitchev A. Repression of the human dihydrofolate reductase gene by a non-coding interfering transcript. Nature. 2007 Feb;445(7128):666–70.

67. Costello JT, Bieuzen F, Bleakley CM. Where are all the female participants in Sports and Exercise Medicine research? European Journal of Sport Science. 2014 Nov;14(8):847–51.

68. Bickel CS, Slade J, Mahoney E, Haddad F, Dudley GA, Adams GR. Time course of molecular responses of human skeletal muscle to acute bouts of resistance exercise. Journal of Applied Physiology. 2005 Feb;98(2):482–8.

69. Yang Y, Creer A, Jemiolo B, Trappe S. Time course of myogenic and metabolic gene expression in response to acute exercise in human skeletal muscle. Journal of Applied Physiology. 2005 May;98(5):1745–52.

70. Louis E, Raue U, Yang Y, Jemiolo B, Trappe S. Time course of proteolytic, cytokine, and myostatin gene expression after acute exercise in human skeletal muscle. Journal of Applied Physiology. 2007 Nov;103(5):1744–51.

71. Shi H, Zhou Y, Jia E, Pan M, Bai Y, Ge Q. Bias in RNA-seq Library Preparation: Current Challenges and Solutions. Biomed Res Int. 2021;2021:6647597.

72. Khrameeva EE, Gelfand MS. Biases in read coverage demonstrated by interlaboratory and interplatform comparison of 117 mRNA and genome sequencing experiments. BMC Bioinformatics. 2012 Apr 19;13 Suppl 6(Suppl 6):S4.

73. Lahens NF, Kavakli IH, Zhang R, Hayer K, Black MB, Dueck H, et al. IVT-seq reveals extreme bias in RNA sequencing. Genome Biol. 2014 June 30;15(6):R86.

74. Kukurba KR, Montgomery SB. RNA Sequencing and Analysis. Cold Spring Harb Protoc. 2015 Apr 13;2015(11):951–69.

75. Illumina Knowledge [Internet]. [cited 2025 July 11]. Available from: https://knowledge.illumina.com/~gitbook/pdf?page=wNfzOcuMnPCFZ2eulp1Q&only=yes&limit=100

76. Wilusz JE. Long noncoding RNAs: Re-writing dogmas of RNA processing and stability. Biochim Biophys Acta. 2016 Jan;1859(1):128–38.

77. Telzrow CL, Zwack PJ, Esher Righi S, Dietrich FS, Chan C, Owzar K, et al. Comparative analysis of RNA enrichment methods for preparation of Cryptococcus neoformans RNA sequencing libraries. G3 (Bethesda). 2021 Oct 19;11(11):jkab301.

